# Metatranscriptomics reveals declines in ice cover influence winter viral community activity

**DOI:** 10.1101/2024.04.01.587580

**Authors:** Elizabeth R. Denison, Brittany N. Zepernick, R. Michael L. McKay, Steven W. Wilhelm

**Affiliations:** Department of Microbiology, University of Tennessee, Knoxville, Tennessee, USA; Great Lakes Institute for Environmental Research, University of Windsor, Windsor, Ontario, Canada

**Author notes:** Correspondence: Steven W. Wilhelm, 1-865-974-0665.

**Keywords:** Viral ecology, winter limnology, metatranscriptomics, climate change, diatom bloom

## Abstract

Freshwater lakes are sentinels of environmental change, and climate change-driven declines in ice cover have been shown to disrupt aquatic communities and jeopardize ecosystem services. Viruses shape microbial communities and regulate biogeochemical cycles by acting as top-down controls, yet there is relatively little known about how declining ice cover will influence viral community activity. Lake Erie is a critical freshwater ecosystem and serves as a model system to assess how ice cover extent will affect winter limnology. We surveyed size selected surface water metatranscriptomes for conserved viral hallmark genes as a proxy for active virus populations and compared activity profiles between ice-covered and ice-free conditions from two contrasting winters. Active virus communities were present in both conditions, spanning diverse phylogenetic clades of bacteriophage (*Caudovirales*), giant viruses (*Nucleocytoviricota*), and RNA viruses (*Orthornavirae*). However, viral activity was significantly shaped by the extent of ice cover. Notably, viral richness and relative transcript abundance in the surface waters were reduced under ice relative to the ice-free conditions. Correlations with microbial community metrics suggest the differences in viral communities are at least in part driven by the decreased winter diatom bloom associated with declines in ice cover. Overall, our data suggest viral community activity is influenced by ice cover extent, and viruses may serve as sentinels of environmental disturbance and ecosystem response(s) to climate change.

**IMPORTANCE:** As ice cover is projected to become increasingly rare on large temperate lakes, there is a need to understand how microbial communities during winter months might respond to these changing ice-cover conditions. Despite the documented controls viruses have on microbial communities, little is known regarding the relationship between virus activity and ice cover extent. By using metatranscriptomics to investigate virus communities, we show that viral community activity is sensitive to ice cover extent, likely due in part to ice cover-driven shifts in host community structure. This work serves to build our understanding of how viral communities will function in a future, potentially ice-free, climate.

## INTRODUCTION

Decades of research have demonstrated viruses are key biological components of aquatic ecosystems (1). As top-down controls, viruses influence microbial community structure (2) and the movement of organic matter *via* the “viral shunt” and “viral shuttle” (3, 4). Viruses also modulate the metabolism of infected cells by hijacking host cell physiology and through virus-encoded metabolic genes (5), which can have far-reaching implications on biogeochemical cycles (6–9). As many of Earth’s aquatic ecosystems are faced with unprecedent changes in climate (10, 11), there is growing interest in how viral communities are influenced by or amplify climate change and in their incorporation into predictive models (12). Temperate fresh waters are particularly vulnerable to climate change (13–15), where one manifestation is changes in ice phenology (16). In fact, the frequency of ice-free winters is projected to increase globally in current climate trajectories (17). Seasonally ice-covered lakes are not biologically dormant during the winter (18–20), and yet there is relatively little known about viral activity during this period (21).

Lake Erie is a seasonally ice-covered North American Great Lake that supports key socioeconomic and ecosystem services, many of which are threatened by human disturbances and climate change (22–25). Winter in the Laurentian Great Lakes is undergoing pronounced changes due to warming climate (16), and since the 1970’s Lake Erie has exhibited significant declines in ice cover extent and duration (26, 27). Furthermore, ice cover extent drastically alters microbial community structure by shifting the winter ecosystem from one characterized by dense accumulations of filamentous diatom biomass (28–30) to that of picoplankton (31). The response of viral communities to declining ice cover, however, has not been explored to our knowledge. Given that viruses are entangled with carbon and nutrients cycles as well as food web dynamics in Lake Erie (32–34), it is key to understand how declining ice cover may impact viral community activity.

Viruses have been detected during mid-winter in Lake Erie (35, 36), but there has been no comprehensive characterization of winter virus activity in the Great Lakes. Viruses are likely affected by ice cover extent as ice can shape host population structure (37–39) in addition to there being other ice cover-dependent factors (*e*.*g*., UV exposure and decay rates (40, 41)) that are intrinsically linked to viral activity as well. Lakes are sentinels of climate change (13), and Lake Erie represents a unique model environment for investigating how climate change manifests through ice cover extent. Microbial communities also act as sentinels (42) and understanding viral community dynamics may help reveal the downstream effects of declining ice cover on aquatic ecosystems.

Targeted gene amplification and shotgun metagenomic approaches have been informative in the study of viruses in the Great Lakes (35, 43–45), but a limitation of DNA-based approaches is they do not reveal virus activity (replication) or capture RNA viruses. Metatranscriptomics can facilitate both the discovery of RNA viruses as well as estimate *in situ* activity levels of viruses using transcript abundance as a proxy. Here, we surveyed metatranscriptomes from size-selected (> 0.2 µm) samples collected from Lake Erie surface waters during two contrasting winters: one with high-ice cover (2019, 95% mean maximum ice cover) and a subsequent relatively ice-free winter (2020, 15% mean maximum ice cover) (46). Samples from the spring months following the ice-free winter were also collected and serve as an outgroup in this study. Using hallmark genes as proxies for viral populations, we identified phylogenetically diverse viral communities active in the winter water column. Active viral community structure and composition were shaped by ice cover extent, wherein viral richness and relative transcript abundance was reduced under ice compared to the ice-free conditions. While the drivers behind the ice cover-associated differences in viral communities are unclear, viral richness and abundance were correlated with corresponding differences in microbial community structure and diversity.

## MATERIALS AND METHODS

### Sample collection, RNA extraction, and sequence processing

Opportunistic surface water samples were collected by USCGC *Neah Bay* between February and March of two consecutive years (2019 and 2020) (47). The winter of 2019 was considered a high-ice year (95% mean maximum ice cover) while the winter of 2020 was largely ice-free (15% mean maximum ice cover) (46). Additional spring samples in 2020 were collected by MV *Orange Apex* (47). Samples were collected from 0.5 meters below the surface as either whole-water (n = 20) or plankton-net concentrated (n = 57). In this work, we examined the whole-water samples to account for the effect of collection method on the metatranscriptome profiles (Figure S1). Please refer to the Supplemental Material for details regarding collection method. Whole-water samples spanned three conditions: ice-covered (Feb/Mar 2019, n=4), ice-free (Feb/Mar 2020, n=10), and spring (May/Jun 2020, n=6). Samples intended for RNA extraction were passed through 0.22-µm nominal pore-size filters, flash-frozen, and stored at -80 °C until further processing. Details regarding RNA extraction and sequence processing can be found in an associated *Microbiology Resource Announcement* (47). Sample metadata can be accessed through the Biological and Chemical Oceanography Data Management Office (BCO-DMO) under dataset number 809945 (48).

### Metatranscriptome assembly and gene prediction

This study used the metatranscriptome co-assembly and gene predictions generated by Zepernick *et al*. (49). Briefly, in Zepernick *et al*., quality-filtered reads were co-assembled using MEGAHIT v1.2.9 (50) and the resulting co-assembly was assessed using QUAST v5.0.2 (Table S1) (51). Gene sequences were predicted from the co-assembly contigs using MetaGeneMark v3.38 (52). Here, to quantify the relative transcript abundance for genes within a sample we mapped trimmed reads to gene sequences with ≥ 90% identity and ≥ 90% read length fraction thresholds (53) using CoverM v0.6.1 (54). We defined a gene as “present” in a metatranscriptome library if at least 50% of the gene was covered (55). Read counts for genes with less than 50% coverage were reset to zero on a per-library basis. Read counts were converted to transcripts per million (TPM) (56).

### Viral hallmark gene discovery

Conserved viral hallmark genes were identified as proxies for viral populations to estimate viral diversity and activity, similar to as performed previously (57, 58). Here, viral hallmark genes detected in the metatranscriptomes were interpreted to represent “active” or cell-associated viral populations. This was presumed because 1) DNA viruses must be replicating intracellularly to synthesize mRNA and 2) samples were filtered prior to nucleic acid extraction at a size cutoff that would enrich for cell-associated RNA viruses (> 0.2 µm).

All hallmark genes were identified through protein sequence similarity searches. Predicted protein sequences from MetaGeneMark were aligned to databases of viral hallmark genes (detailed below) using DIAMOND BLASTP (59) (E-value threshold of 1e^-5^). Protein sequences that aligned to translated hallmark genes were then aligned to the RefSeq v213 database and sequences with a top hit to a viral protein were retained as putatively viral (58, 60). As a second quality control measure, only viral sequences with an appropriate protein domain identified by a Pfam domain search (database v32) were retained for analysis (*i.e.,* a putative Gp23 sequence retrieved from the BLASTp that had no capsid domain was not included). RdRp-like protein sequences were further filtered using Palmscan (-rdrp option) and only high confidence RdRp sequences retained (61). As an additional measure to remove non-viral sequences, the metatranscriptome contig from which the hallmark gene originated was piped through VirSorter2 v2.2.3 (62) and CheckV v1.0.1 (63). While the majority of contigs were too short to determine if they were non-viral, no hallmark gene contigs had any cellular genes called by VirSorter2 or CheckV.

### Hallmark gene databases

The terminase large subunit (TerL) was used as a general phage marker (*i.e.,* a marker that is present in all *Caudovirales*) (Table S2) (64). Relatively few TerL were detected, most of which resembled T4-like myophage based on sequence homology. Therefore, the T4-like major capsid protein (Gp23), DNA polymerase B (Gp43), and tail sheath domains were used to detect myophage. The Gp23 had the greatest richness compared to the other phage markers and was therefore chosen as the representative hallmark gene for myophage for further analysis (Table S2). Viral integrase, excisionase, CI repressor, and Cro repressor were also screened for as these markers have been used to detect lysogenic phage in metagenomic data (65, 66). All *Caudovirales* marker sequences were retrieved from Interpro, where only viral sequences were downloaded.

The major capsid protein (MCP) was used as a proxy for active *Nucleocytoviricota*, or NCLDV, populations. We chose the MCP marker because 1) it is conserved in many NCLDV genomes (67) (all known groups except for *Pandoraviridae* (68) and *Pithoviridae* (69)), and 2) it is most likely to be detectable since the MCP can be highly expressed relative to other NCLDV core genes (58, 70). Because the MCP may not be preferred for phylogenetics (71), we also searched for DNA polymerase B (PolB) as an alternative NCLDV marker. Only nine PolB sequences were detected in the metatranscriptomes *via* read mapping (Table S2). Hereafter, we only discuss the MCP as a proxy for NCLDV with the caveat that our phylogeny is an estimate. NCLDV marker sequences were retrieved from the NCVOG database (72).

RNA-dependent RNA polymerase (RdRp) was used as the hallmark gene for RNA viruses (*Orthornavirae*) (73), and the RdRp-Scan (74) database was used for RdRp searches. Replicase sequences compiled by Kazlauskas *et al*. (75) and Moniruzzaman *et al*. (58) were used to screen for circular replication-associated protein-encoding single-stranded (CRESS) DNA viruses (*Cressdnaviricota*).

### Phylogenetic analysis

Hallmark protein sequences were aligned to reference alignments using Clustal Omega v1.2.4 (76). Positions with >90% gaps were trimmed from the alignments by trimAl v1.4 (77). Trees were constructed using FastTree v2.1.11 (-lg option) (78) or IQ-TREE v2.2.0.3 (-m TEST option) (79) and were visualized in iTOL (80). NCLDV reference genomes were retrieved from Gilbert *et al*. (81) and the MCP sequences were predicted using ViralRecall v2 (82). RdRp-Scan alignments were used for the RdRp tree references (74). References for all other trees were manually curated.

### Microbial community characterization

Taxonomic and functional annotation of predicted cellular genes were retrieved from Zepernick *et al*. (49). Briefly, taxonomy was estimated using EUKulele v1.0.6 (83), where protein sequences were aligned to the PhyloDB database v1.076 by DIAMOND BLASTp. Cellular genes were functionally annotated by eggNOG-mapper v2.1.6 using the eggNOG 5.0 orthology database with a DIAMOND BLASTp cutoff of 1e^-10^ (84, 85).

### Statistical analyses

Community diversity metrics were calculated using the R package vegan (86, 87) and in PRIMER v7 (88). Normalized transcript abundance (TPM) tables were used as the input for all diversity analyses. For beta diversity statistics, Bray-Curtis dissimilarity matrices were generated from the standardized TPM table (Wisconsin double standardization). Similarity percentage (SIMPER) analyses were used to quantify the contribution of individual marker genes to average dissimilarity between sample groups.

Spearman’s correlations were performed using R (cor.test) (86). The virus-host hallmark gene interaction network was generated using FlashWeave (*α* < 0.01) (89), where genes annotated as RNA polymerase subunits Rpb1 (KEGG orthology K03006) and RpoB (K03043) were used as cellular hallmark genes (53, 90, 91). A TPM table of all viral and cellular hallmark genes was used as input.

### Data availability

Raw and quality-filtered reads are publicly available through the JGI Genomes Online Database (GOLD) under GOLD Study ID Gs0142002.

## RESULTS

### Signatures of active viral communities within the winter surface water metatranscriptomes

Viruses were active in the winter surface water samples, with all viral groups detected by our hallmark gene search (Figure 1A). Overall, active winter viral communities displayed high hallmark gene richness and were phylogenetically diverse (Figure 1A-D). The majority of Gp23 detected in the winter metatranscriptomes via read mapping (439 predicted gene sequences) were most closely related to uncultured myophage environmental sequences with uncertain hosts as opposed to the established cultured clades, although Gp23 were related to isolated cyanophage (two sequences) and *Pelagibacter* phage (six sequences) (Figure 1B, Table S5). The NCLDV MCP detected in the winter libraries (427 sequences) represented five of the six established NCLDV orders (Figure 1C) (71), most of which were assigned to the *Imitervirales* (395 sequences) (Table S3). The relatively higher *Imitervirales* MCP richness may reflect the broad host range of this group (92, 93). The phylogenetic diversity of RdRp present in the winter libraries (332 sequences) was similarly high, spanning the known phylum-level diversity of the *Orthornavirae* (Figure 1D, Table S3). Furthermore, only a single lysogenic marker (integrase) was considered viral based on its top BLASTp hit to uncultured fresh water *Caudovirales* (CAB5220699.1) and was only present in one library in the ice-free conditions (MT6, 02/14/2020).

**Figure 1.**
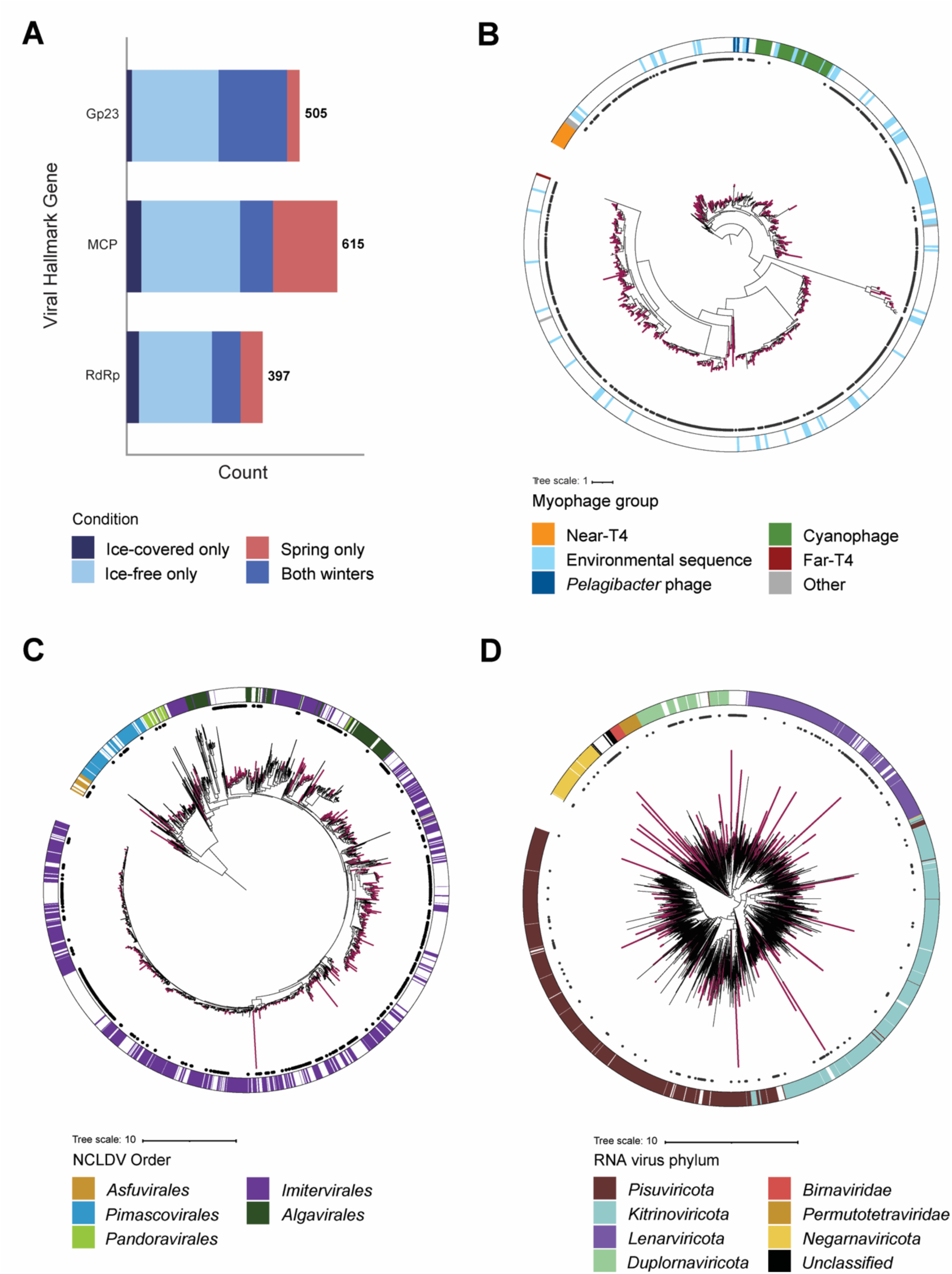
Viral hallmark genes detected in the surface water metatranscriptomes. **A)** Total count of genes discovered for Gp23, MCP, and RdRp. Maximum-likelihood trees for **B)** Gp23, **C)** MCP, and **D)** RdRp. Branches for Lake Erie sequences are bolded and in color, reference sequence branches are in black. Inner ring circles represent Lake Erie sequences detected in at least one winter library. Outer ring shows taxonomy of reference sequence.

The viral hallmark gene transcript pool was consistently dominated by Gp23 and RdRp transcripts across both winters, although MCP representation increased in the spring, specifically the May 1-22 samples (Figure S2). Only eight single-stranded DNA virus replicase markers were detected with low representation in the water column libraries (Table S2, Figure S2) and they were not analyzed further.

### Viral community activity profiles are distinct between the ice-covered and ice-free conditions

Using the relative transcript abundance of the viral hallmark genes Gp23, MCP, and RdRp as a proxy for viral community activity revealed activity profiles between the ice-covered and ice-free conditions were highly dissimilar (73% dissimilar on average) and significantly different between ice cover states (ANOSIM R = 0.79, p = 0.003) (Figure 2A). Furthermore, ice cover extent shaped active viral community composition, meaning the presence or absence of viral hallmark genes (ANOSIM R = 0.86, p = 0.002) (Figure 2B). Mean viral community evenness (Pielou’s) and diversity (Shannon’s H) were not significantly different between ice cover conditions (Figure 2C, Table S4). This is perhaps because most individual hallmark genes had consistently low relative abundance across samples, resulting in high sample evenness. Mean hallmark richness, however, was significantly lower (Tukey HSD p < 0.001) in the ice-covered condition compared to the ice-free (Figure 2C). In other words, there were significantly fewer viral hallmark genes detected via read mapping in the ice-covered winter samples. Viral hallmark richness was still significantly lower when normalized to library size (Tukey HSD p < 0.001).

**Figure 2.**
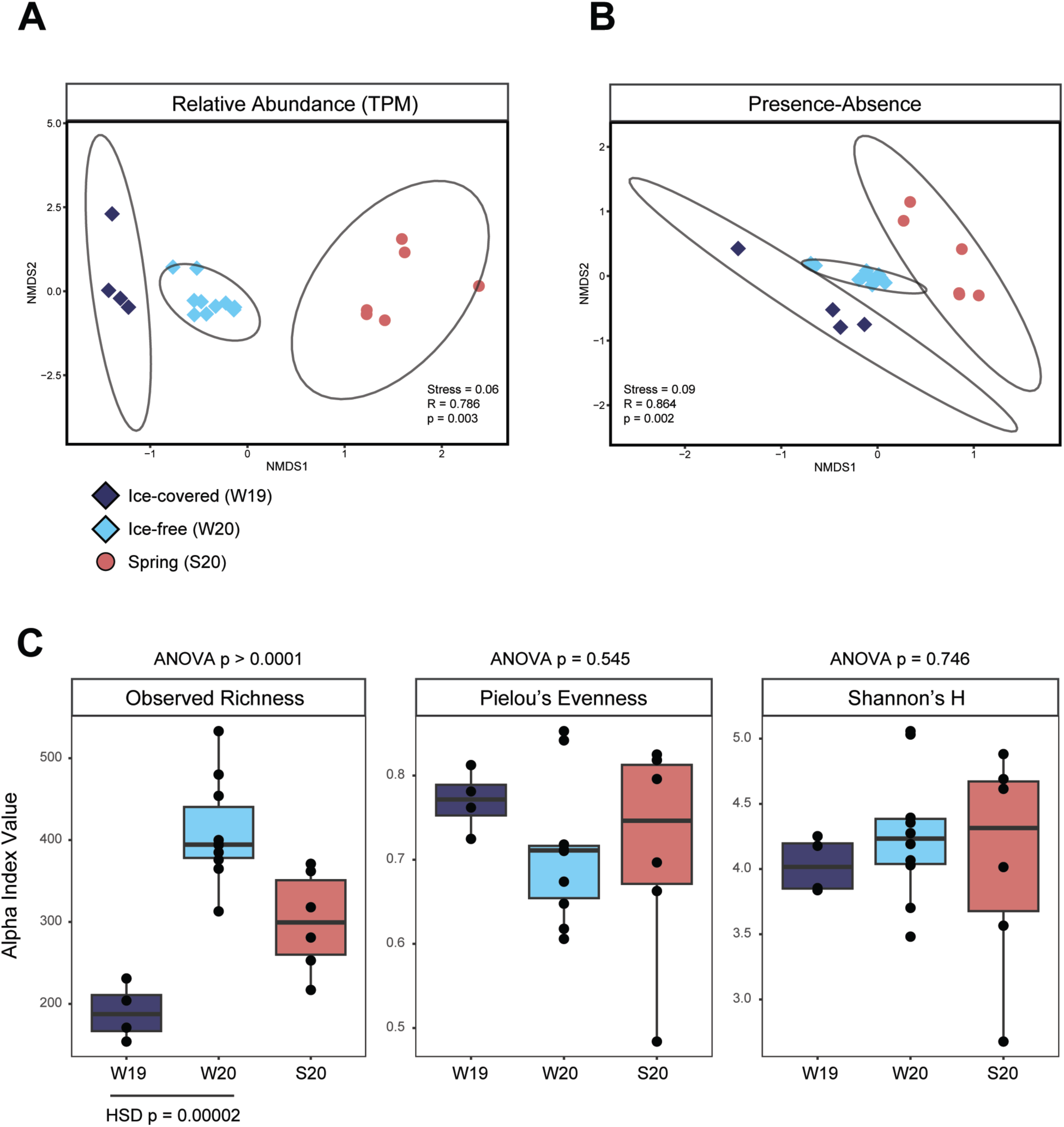
Diversity and composition of viral activity profiles in the surface water metatranscriptomes. Beta-diversity non-metric multidimensional scaling (nMDS) plot illustrating similarity of sample structure based on **A)** relative transcript abundance and **B)** presence-absence transformed abundance of viral markers. Ellipses represent 95% confidence intervals for the three seasons sampled. **C)** Alpha diversity metrics grouped by season. Tukey HSD is shown for the ice-covered winter 2019 (W19) and ice-free winter 2020 (W20) sample comparison when applicable.

Viral community activity was significantly different between winters when examining virus hallmark types individually. The separate Gp23, MCP, and RdRp activity profiles all clustered by season and were significantly different between the ice-covered and ice-free winters (Figure S3). Similarly, the mean richness of all three virus types was significantly lower in the ice-covered winter relative to the ice free (Figure S4, Table S5). Overall, active viral communities are shaped by ice cover extent (ice-covered or ice-free conditions) in regard to active viral community composition and structure, and ice cover influenced a broad range of virus types.

### Ice-associated shifts in viral activity profiles are associated with differences in microbial community activity

Collectively, viral hallmark genes comprised a larger portion of the ice-free communities (*i.e.,* total TPM) compared to the ice-covered (Figure 3A). Higher viral abundance in the ice-free conditions coincided with lower abundance of the bloom-forming diatoms (*Bacillariophyta*), both in terms of lower diatom transcript abundance (Figure 3B) and cell counts (49). Conversely, prokaryotic transcripts had higher representation in the ice-free conditions (13-36% versus 30-67%, respectively) (Figure 3B). In fact, collective prokaryotic normalized abundance was significantly negatively correlated with diatom normalized abundance across the winter samples (Spearman’s ρ = -0.83, p < 0.001) (Table S6). Moreover, both prokaryotic (*rpoB*) and eukaryotic (*rpb1*) hallmark gene richness was lower in the ice-covered winter (Figure S5). The ice cover-associated shifts in microbial community structure were correlated with the observed differences in viral abundance and richness. Notably, viral hallmark normalized transcript abundance was significantly negatively correlated with diatom transcript abundance (ρ = -0.82, p < 0.001) (Figure 3C) and positively so with prokaryotic abundance (Table S6). Furthermore, viral richness was positively correlated with microbial community richness (RpoB/Rpb1) (ρ = 0.93, p = 2.2e-16) (Figure 3D). Altogether, lower diatom representation in the ice-free metatranscriptomes was associated with higher microbial community richness and prokaryotic representation, and in turn higher viral richness and representation.

**Figure 3.**
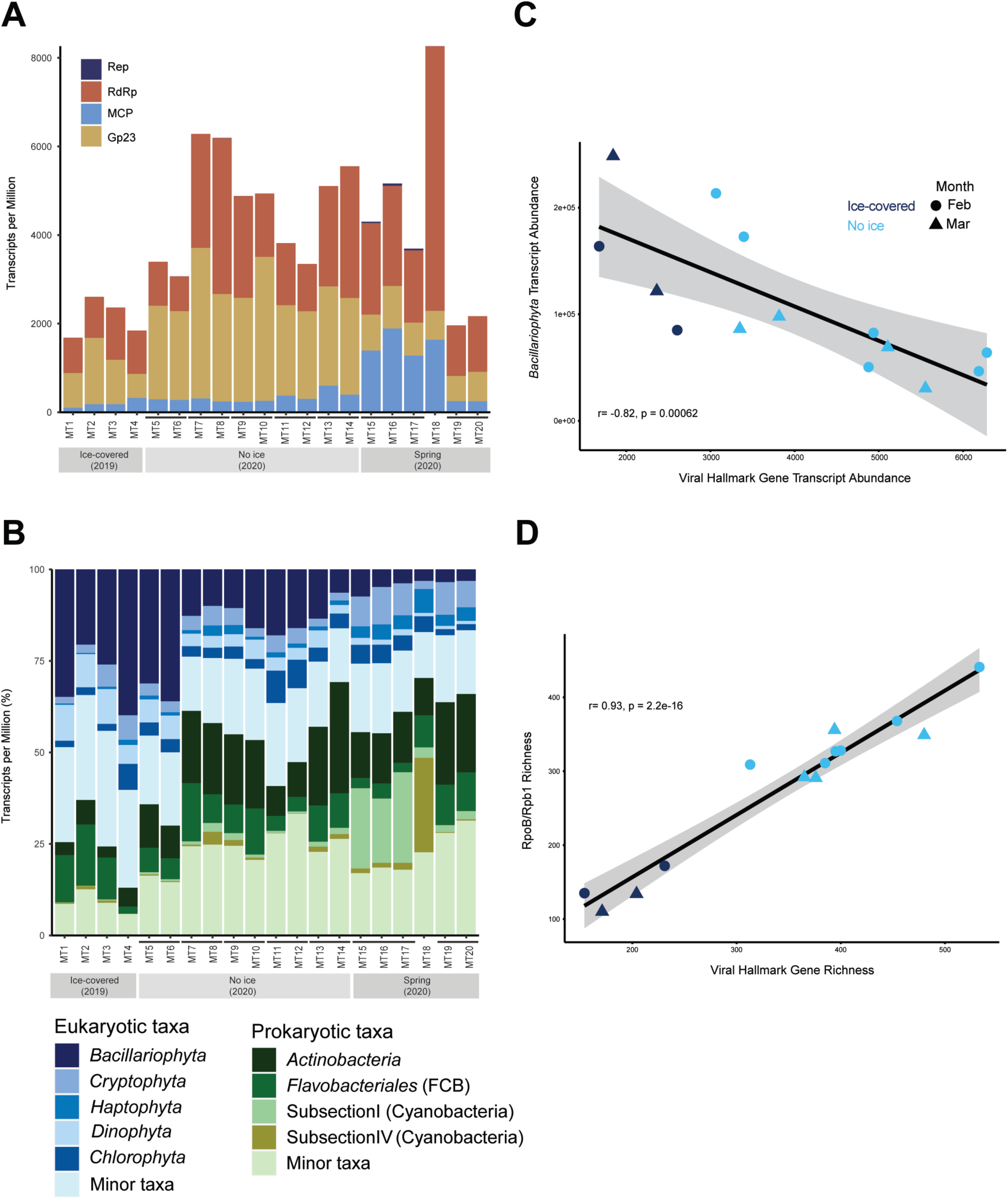
**A)** Relative transcript abundance (TPM) of the viral hallmark genes for myophage (Gp23), NCLDV (MCP), RNA viruses (RdRp) and CRESS DNA viruses (Rep). **B)** Relative transcript abundance of the dominant eukaryotic and prokaryotic taxa. Spearman’s correlations between **C)** viral hallmark and diatom (*Bacillariophyta*) transcript abundance and **D)** viral and cellular hallmark gene richness. Biological replicates are connected by horizontal bars.

### Hallmark genes with the largest contributions to dissimilarity between winter conditions

Although viral richness and representation (collective TPM) were higher in the ice-free conditions, only a select few individual viral markers notably increased and contributed to the dissimilarity between the ice cover conditions. Out of the total 1,525 viral markers, only 11 had greater that 0.5% contribution to average dissimilarity and amounted to the cumulative 10% average dissimilarity between the ice cover conditions (Table S7). While this cut-off is somewhat arbitrary, it captures the viral markers with the largest shifts in relative abundance between the ice cover states, and the remaining markers have low individual contributions (Table S7). Eight of the eleven markers of interest (four Gp23 and four RdRp) had higher average representation in the ice-free relative to the ice-covered winter where they “spiked” in transcript abundance in the ice-free conditions, while the remaining three (one Gp23 and two RdRp) had the opposite trend (Figure 4). Notably, the three markers that had higher average representation in the ice-covered winter had an overall low relative abundance (Figure 4), which is in line with the overall low viral TPM in the ice-covered winter samples noted previously.

**Figure 4.**
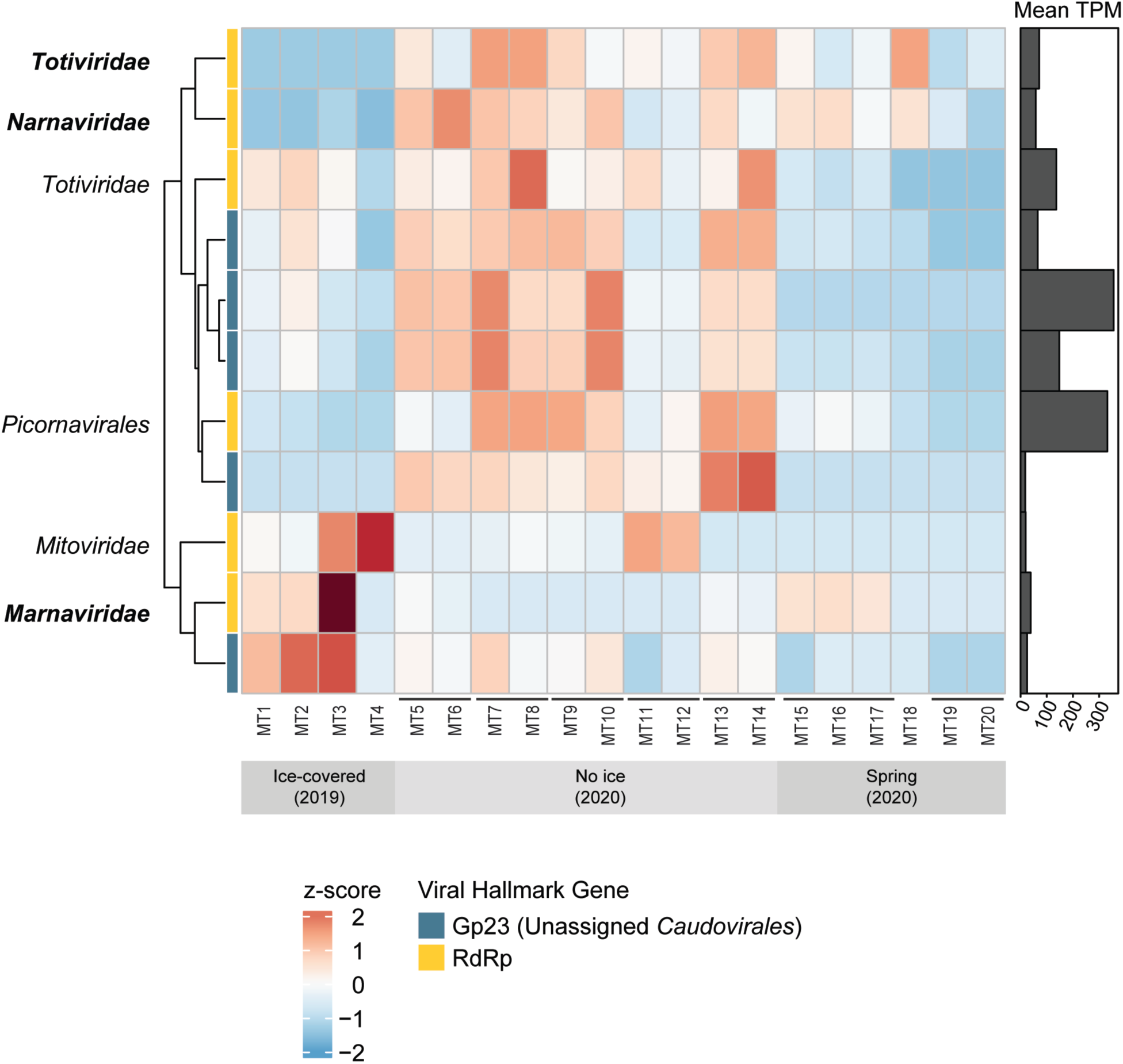
Relative transcript abundance (z-scored TPM) of the eleven top contributors to dissimilarity between the ice-covered and ice-free samples based on SIMPER analysis. Rows are clustered by abundance pattern. Bar plot shows mean TPM across all twenty samples. Biological replicates are connected by horizontal bars. Phylogenetic placement for RdRp are indicated by text labels and diatom-associated RdRp clades are in bold.

All five of the top Gp23 contributors were placed within a clade containing an uncultured freshwater Gp23 sequence with uncertain host range (CAB5226458.1). The RdRp of interest spanned three RNA virus phyla (Figure 4). Interestingly, three RdRp were related to diatom viruses within the *Bacillarnaviridae* (order *Picornavirales*) (Figure S6) and diatom colony associated viruses within the *Narnaviridae* (phylum *Lenarviricota*) (Figure S7) and *Totiviridae* (phylum *Duplornaviricota*) (Figure S8). To further infer virus-host pairs, we assessed correlations in relative transcript abundance between the cellular marker genes (*rpoB*/*rpb1*) and the virus hallmark genes, but no significant interactions (*α* < 0.01) were found between the diatom Rpb1 and putative diatom virus RdRp markers.

Major capsid protein transcript abundance was consistently low throughout both winters, and subsequently no single MCP marker had a large contribution to the dissimilarity between winters. However, MCP had higher representation in the spring samples (May 1-22) relative to both winters largely due to the relative increase in two MCP. Their dominance was also likely responsible for the low MCP evenness and diversity in the spring samples (Figure S4). Although host range is uncertain, these MCP resembled isolated Chloroviruses (order *Algavirales*) that infect green algae and *Mimiviridae* (order *Imitervirales*) isolates that infect amoeba (Figure 1C). It was noteworthy that the Chlorovirus-like MCP had a significant positive correlation with a Rpb1 annotated as Chlorophyta (*Trebouxiophyceae*) (*α* < 0.01, cor > 0.8).

### Identification of NCLDV bacteriorhodopsins

Light availability has been implicated as a driving factor behind how declining ice cover shapes microbial communities (94), specifically regarding phytoplankton ecology in Lake Erie (31, 49). Furthermore, viruses can encode functional metabolic genes involved in phototrophy and light-sensing, namely the D1 protein (PsbA) in cyanophage (95) and rhodopsin in NCLDV (96). We identified three putative NCLDV bacteriorhodopsins (Figure S9), indicating actively transcribed viral rhodopsins in Lake Erie. However, no trends in transcript abundance between the ice-covered and ice-free conditions emerged. Both homologues were only detected in the spring samples, which was expected as NCLDV MCP transcript abundance was low throughout both winters (Figure 3A). For further details on this analysis, please refer to the Supplemental Material.

## DISCUSSION

Hallmark genes for diverse viruses were detected in the winter metatranscriptome libraries, indicating an “active” virus community present during winter in Lake Erie. Whereas virus-like particles and sequences have been enumerated from Lake Erie winter samples before (35, 36), this study greatly expands the known diversity of active viruses. Although phylogenetically diverse, most individual hallmark genes had low relative transcript abundance within a library, and the viral hallmark transcript pool was disproportionally dominated by a small fraction of the total intra-sample richness. This suggests a community functioning in a manner similar to the Bank model proposed by Brietbart and Rohwer (97). The Bank model proposes local virus diversity is often high, but most viruses are rare and inactive and that only a small fraction of viruses with sufficient susceptible host populations are replicating (97). Although in this study we cannot infer the percentage of viruses that are active (*i.e.,* we could not enumerate free virus particles or viral genomes), it appears that within active populations the Bank model is observed.

Ice cover extent shaped active viral community structure and composition. Notably, viral hallmark gene richness and relative abundance was higher in the ice-free condition, largely due to the “spike” in a select few hallmark genes. Our findings suggest the observed differences in active viral communities reflect the ice cover-driven shifts in microbial community structure. Similar to previous observations (31, 49), we observed evidence of a decrease in magnitude of the winter diatom bloom in the ice-free conditions compared to the ice-covered state. It is unclear the degree that host population structure influences winter viral communities, but the differences in host representation and diversity between the ice-covered and ice-free states are in part responsible. Other studies have attributed lower viral abundance or richness in the winter to lower potential host pool abundance (98, 99). Moreover, in principle viral diversity and abundance is shaped by the surrounding susceptible host diversity, productivity, and density (37, 39, 100). Perhaps the increase in prokaryotic representation and cellular marker diversity in the ice-free winter (corresponding to the decrease in magnitude of the winter diatom bloom) promoted the increase in viral richness.

The larger representation of virus activity in the ice-free winter was primarily due to a small fraction of the total marker community. Interestingly, these contained hallmark genes similar to those of diatom-associated RNA viruses, including one RdRp related to lytic *Bacillarnavirus* (order *Marnaviridae*) isolates that infect marine diatoms (101), as well as RNA viruses associated with an environmental diatom colony (102). Culture-dependent studies have shown titers of some diatoms viruses to increase corresponding with the proliferation of diatom blooms and have been speculated to control the bloom crash (103) but may also coexist during the bloom course (104). However, none of the putative diatom RNA viruses interacted with diatom (or any) Rpb1 markers based on our abundance-based network analysis.

We note that active NCLDV did not have large contributions to the differences in viral community activity profiles between ice-cover states, and NCLDV were relatively low in transcript abundance during both winters. Low to undetectable levels of NCLDV (gene copy numbers and transcript markers) during the wintertime have been observed in previous studies surveying the Great Lakes (36, 44, 105). Our findings support the idea that NCLDV activity is generally low during winter months in the Great Lakes, and perhaps NCLDV will be less affected by changing winter ice-cover (at least in terms of activity levels), but more extensive year-round surveys are needed to resolve this. Finally, based on our rhodopsins analysis, NCLDV may have a yet unexplored role in the intersection between ice cover extent, light availability, and phytoplankton physiology. This awaits further study.

We considered the possibility of lysogeny being the predominant phage lifestyle during the winter. To our knowledge no study has specifically investigated lysogeny in Lake Erie, but it has been posited that some phage may “overwinter” as prophage (35). Additionally, other work has suggested ice-cover damage to bacterial cells can promote lysogeny in some scenarios (106). While we do not dismiss lysogeny as a component of Lake Erie microbial communities, we did not observe any clear trends in phage lifestyles between winter climates. In fact, we detected little evidence of lysogenic lifestyles altogether, and lysogeny was not the dominant replication strategy in our surface water winter samples.

### Conclusions

Lakes are sentinels of climate change (13), and this work reveals that active viral communities are sensitive to ice cover decline. Overall, diverse communities of viruses are active in the winter in Lake Erie, but ice-cover extent shapes active viral community profiles as well as the microbial communities with which they interact. As seasonal ice cover is projected to continue to decline (15, 17), winter viral communities are likely to be altered. Perhaps greater richness of RNA viruses and transcriptionally active DNA viruses in ice-free winters and more frequent “spikes” in viral activity will be a shared response. Continued surveillance of freshwater winters (ice-covered and ice-free years) will benefit our understanding of how a changing climate will affect winter viral communities. Additionally, wintertime in seasonal freshwater systems like Lake Erie is not an isolated event and instead is part of an annual cycle (20), and previous reports have shown connectivity between winter ecology and that in the spring/summer and vice versa (18, 107). Therefore, declines in seasonal ice coverage likely not only affect the wintertime communities but also micro- and macro-communities year-round in aquatic systems. Future work should endeavor to sample year-round in order to elucidate not only the effect of ice cover extent on winter viral communities but on the annual cycle overall. As viruses have significant roles in biological communities, understanding alterations in viral activity and host-virus interactions is important, especially in the face of changing climate.

## ACKNOWLEDGEMENTS

This work was supported by National Institutes of Health, NIEHS grant 1P01ES02328939-01, National Science Foundation grant OCE-1840715 (R.M.L.M. and S.W.W.), NSERC grant RGPIN-2019-03943 (R.M.L.M.), funding from the Simons Foundation (735077), funding from the US Department of Energy, Office of Science, Office of Biological and Environmental Research, Genomic Science Program Grant to JPG, under Award Number DE-SC0020362, and JGI project 503851 (S.W.W. and R.M.L.M.). We acknowledge the *Kenneth & Blaire Mossman Endowment* to UTK. The work conducted by the U.S. Department of Energy Joint Genome Institute, a Department of Energy Office of Science User Facility, is supported by the Office of Science of the U.S. Department of Energy under contract DE-AC02-05CH11231. We are thankful for the members of the U.S. Coast Guard Cutter *Neah Bay* and the MV *Orange Apex* along with Dan Peck and Thijs Frenken for assistance with sample collection and processing. We thank Derek Niles of Orange Force Marine Ltd. for coordinating sample collection during spring 2020 in the face of COVID disruptions that prevented conventional surveillance of the Great Lakes by federal agencies. Finally, we thank George Bullerjahn, Christa Pennachhio (DOE JGI), and Justin Chaffin for their collaboration.

## AUTHOR CONTRIBUTIONS

The project was designed by SWW and RMLM. RNA extractions were performed by BNZ. BNZ provided the metatranscriptome assembly and gene predictions. Data analysis, statistics, and interpretation of results were carried out by ERD. Manuscript drafted by ERD. All authors reviewed and edited the final version of the manuscript.

