## Supplemental Tables, Figures and Methods for "Metatranscriptomics reveals declines in ice cover influence winter viral community activity"

**For submission to: mSystems**

### SUPPLEMENTAL MATERIAL

#### Effect of collection method on community metatranscriptome profiles

Collection method shaped the metatranscriptome profiles (Figure S1). Bacterial reads generally had higher representation in the whole-water samples (no filtration) relative to the plankton-net samples (64-μm opening) (Figure S1A). RNA viruses (RdRp) were the dominate virus type in the plankton-net metatranscriptomes, while phage (Gp23) and NCLDV (MCP) were nearly absent relative to the whole-water samples (Figure S1B). Furthermore, viral community activity profiles distinctly clustered by collection method (Figure S1C). Notably, the total number of detected hallmark genes was lower in the plankton-net relative to the whole-water samples (Figure S1D). Plankton-net collection may have resulted in the loss of non-filamentous bacteria and microalgae, which may explain the relative lack of transcripts for phage and NCLDV that infect these groups. Given we were interested in characterizing the total virus community, we examined the unfiltered whole-water samples and plankton-net metatranscriptomes are not discussed here further.

#### Viral auxiliary metabolic genes involved in light-sensing and phototrophy

#### Methods

Protein sequences were aligned to databases of PsbA and bacteriorhodopsin (bR) reference protein sequences using DIAMOND BLASTP (1) (E-value threshold of $1e^{-5}$). PsbA were identified using the PF00124 and MF_01379 profiles downloaded from Interpro. Bacteriorhodopsin were identified using the PF01036 profile and the HMM database by Bulzu *et al*. (2). Only genes retaining the appropriate Pfam domain (*i.e*., PhotoRC or Bac_rhodopsin; Pfam database v32) were retained for further analysis. Phylogenetic trees were constructed in the same manner as described in the Materials and Methods section of the main text. Reference sequences for the bR tree were obtained from Schulz et al. (3). Reference sequences for the PsbA tree were manually curated.

**Results**

Out of the bacteriorhodopsin (bR) genes identified (565 gene sequences), three were considered viral based on phylogenetic placement (Figure S9). Current bR phylogenies display two highly supported viral clades (Viral Group I and Viral Group II) (2, 3). One bR was placed within Viral Group I and one within Viral Group II (Figure S9). Both bR were related to that of freshwater *Mimiviridae*-like MAGs (NCLDV MAG classifications by Schulz et. al (3)). An additional bR fragment was placed within a clade of NCLDV MAGs from aquatic environments (Figure S9), though based on sequence similarity (BLASTP) it resembled cellular bR, suggesting a more recent gene exchange between viral and cellular lineages.

Most Lake Erie PsbA were placed within clades of eukaryotic PsbA (285 out of 494 gene sequences) (Figure S10A). The remainder of sequences (57 sequences) were placed in a clade containing cyanobacterial and cyanophage PsbA. Thirty sequences we considered cyanobacterial as they were placed within one of two clades of freshwater cyanobacteria. Determining the origin of the remaining PsbA (bacterial or phage) was less certain. Eight sequences were most closely related to viral PsbA, but none contained the signature viral motifs documented from cyanophage isolates (4).

The putative eukaryotic *psbA* mRNAs dominated the pool of D1 transcripts (represented as summed TPM per season) (Figure S10B), and as expected diatom-like PsbA (39 sequences) had the largest contribution in both winter climates. Collectively, the clade of cyanobacteria and cyanophage PsbA exhibited low transcript abundance relative to eukaryotes but did make up a larger portion in the spring. This spring increase was driven by two PsbA populations that had notably high summed transcript abundance within the cyanobacteria/cyanophage clade (Figure S10A). Neither were considered viral. One sequence was related to *Aphanizomenon* spp. (*Nostocaceae*) PsbA, while the other sequence most resembled a *Synechococcaceae* metagenome assembled genome (NBQ20673.1).

### SUPPLEMENTAL FIGURES

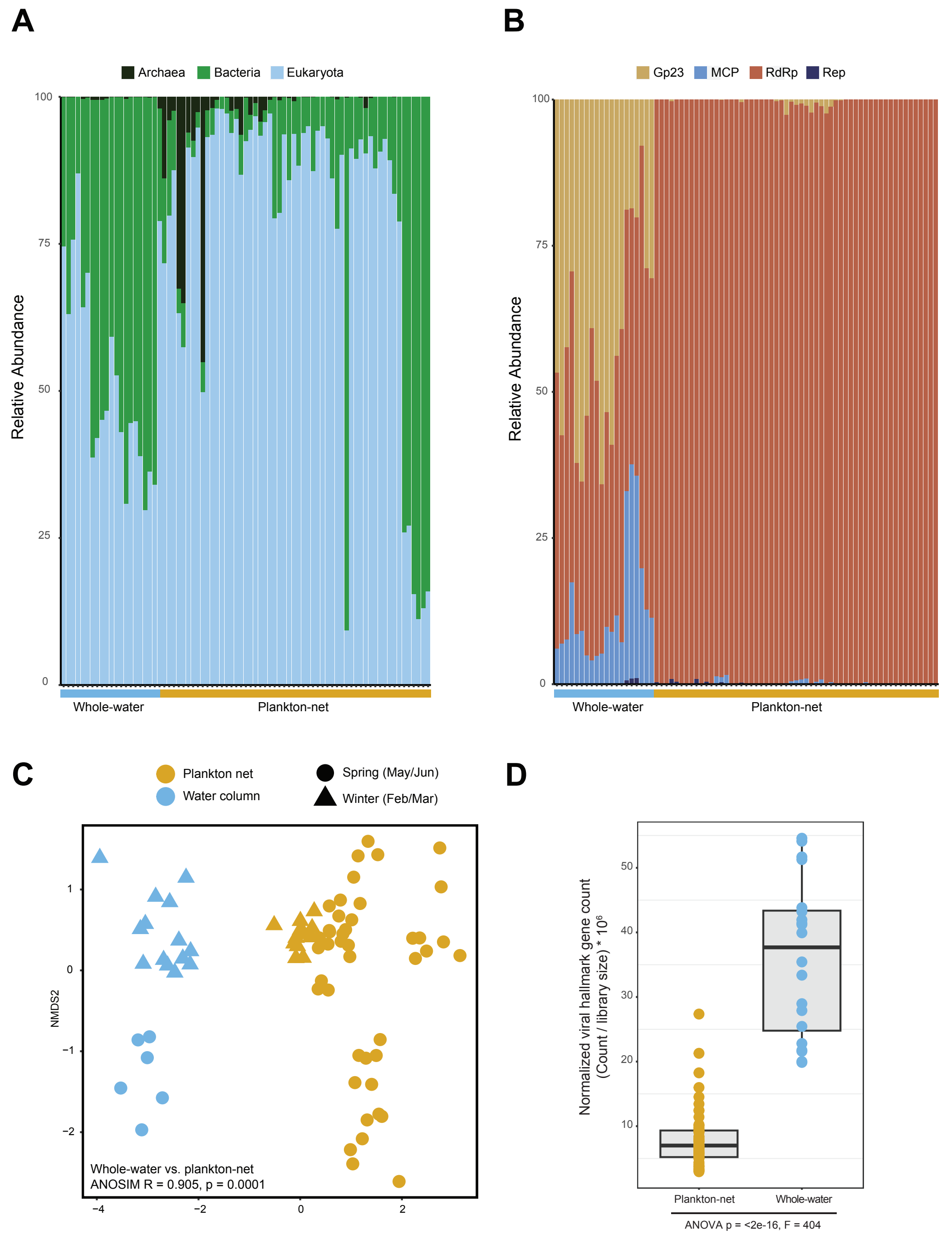

**Supplemental Figure 1.** Influence of collection method on metatranscriptome profiles. Relative abundance (% TPM) of **A)** non-viral predicted genes and **B)** identified viral hallmark genes in the whole-water and plankton-net concentrated samples. **C)** Non-metric multidimensional scaling (nMDS) plot and ANOSIM test results illustrating significant clustering of viral activity profiles by collection method. **D)** Comparison of viral hallmark gene count (observed richness) normalized by library size and scaled.

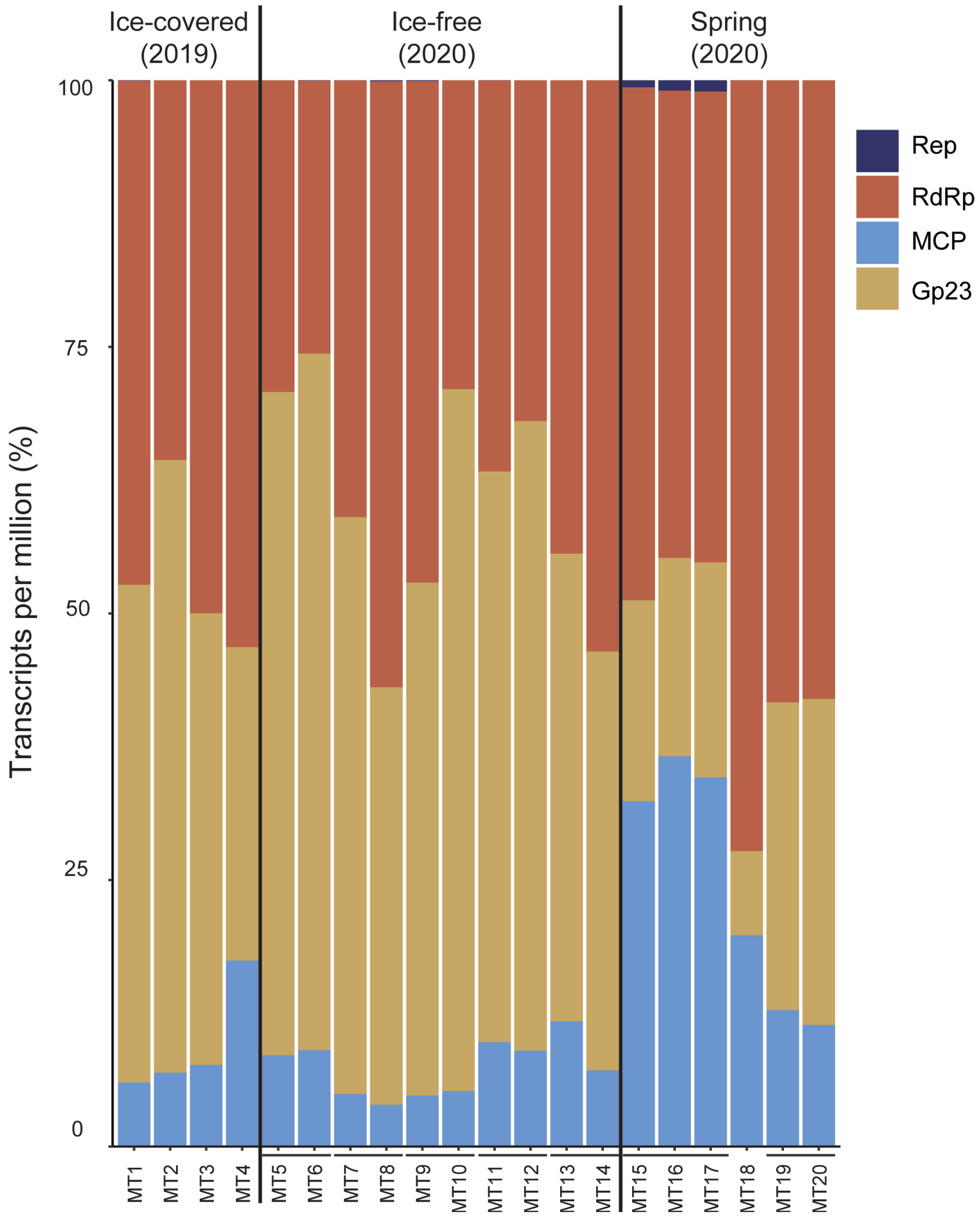

**Supplemental Figure 2.** Relative transcript abundance (TPM) summed by viral hallmark gene type. Biological replicates are connected by horizontal bars.

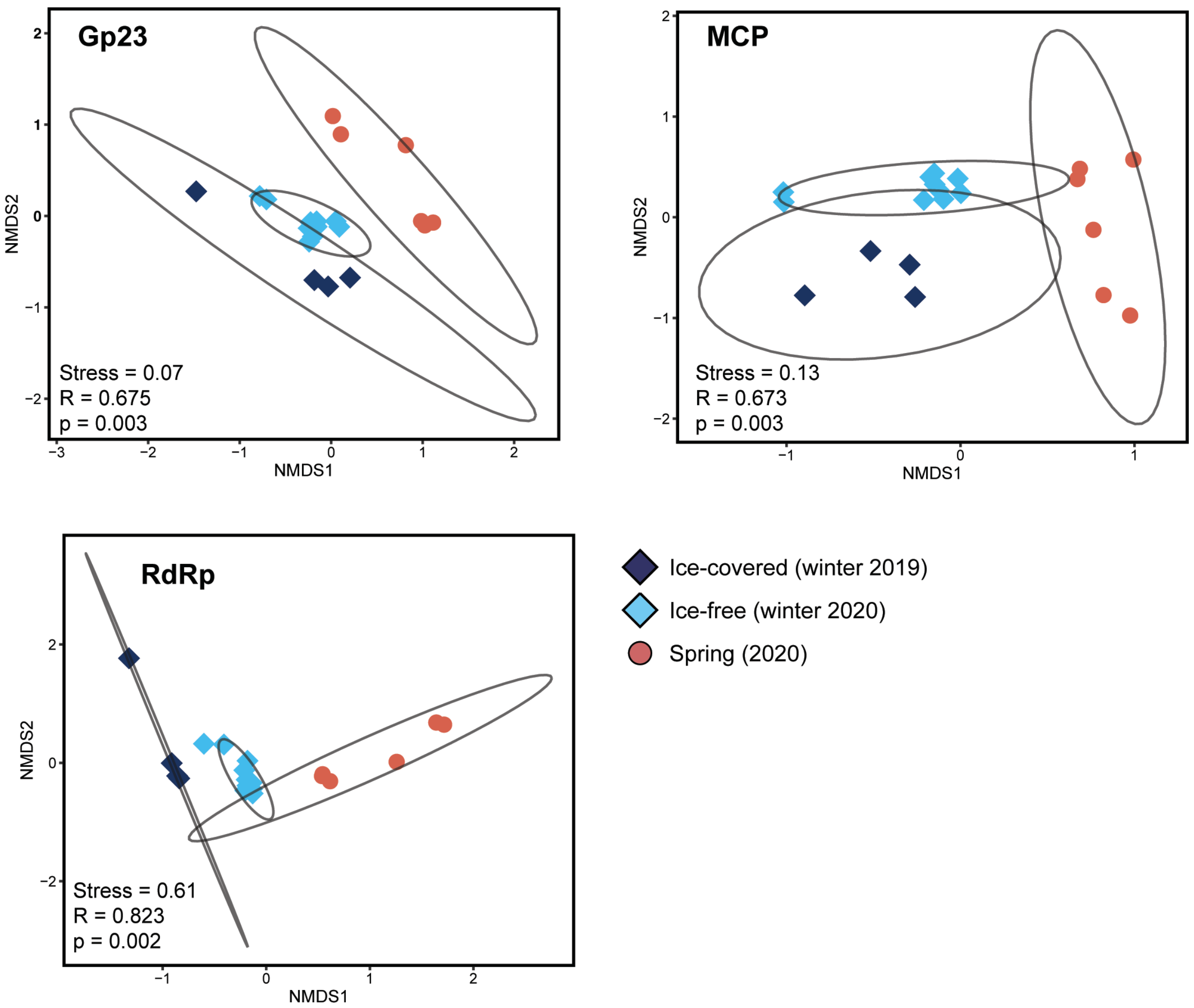

**Supplemental Figure 3.** Beta-diversity non-metric multidimensional scaling (nMDS) plot illustrating clustering of sample structure based on relative transcript abundance (TPM) of individual viral hallmark gene types. Ellipses represent 95% confidence intervals for the three seasons sampled (winter 2019, winter 2020, and spring 2020).

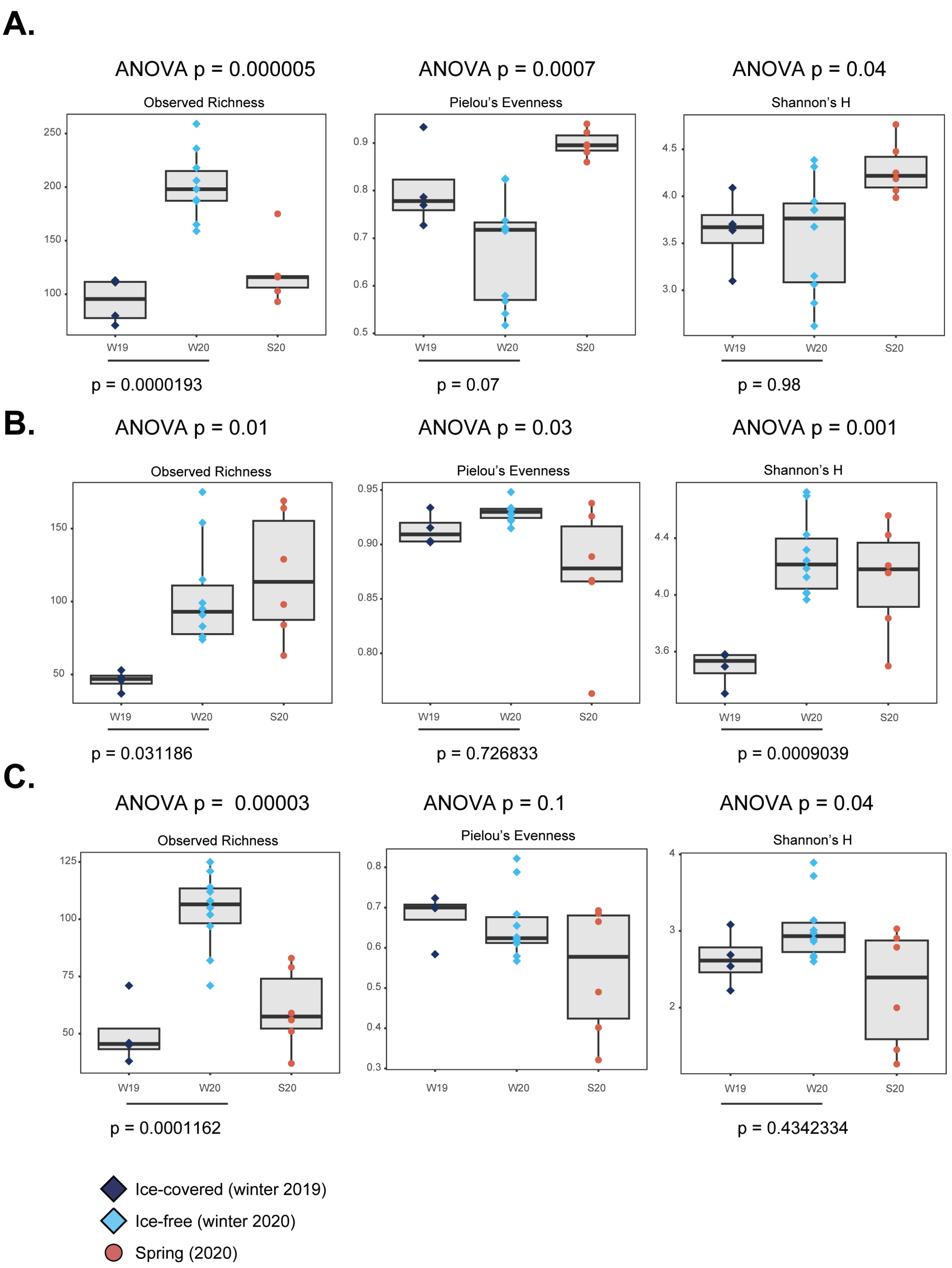

**Supplemental Figure 4.** Alpha diversity metrics grouped by season for the individual viral hallmark gene types **A)** Gp23, **B)** MCP, and **C)** RdRp. Tukey's HSD is shown for the ice-covered (winter 2019, W19) and ice-free (winter 2020, W20) comparison when applicable.

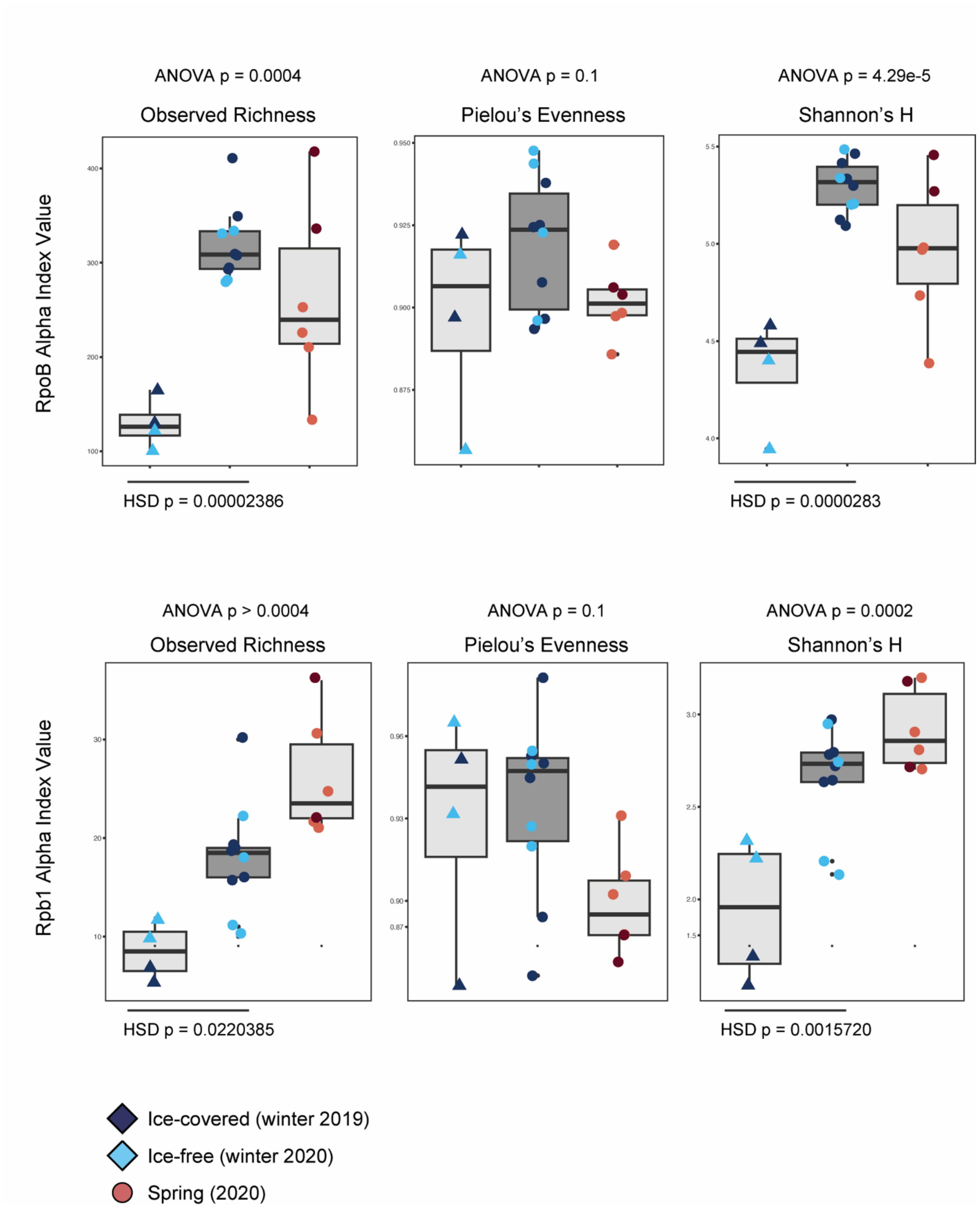

**Supplemental Figure 5.** Alpha diversity metrics grouped by season for RpoB (prokaryotic) and Rpb1 (eukaryotic) marker gene profiles. Tukey's HSD is shown for the ice-covered (winter 2019, W19) and ice-free (winter 2020, W20) comparison when applicable.

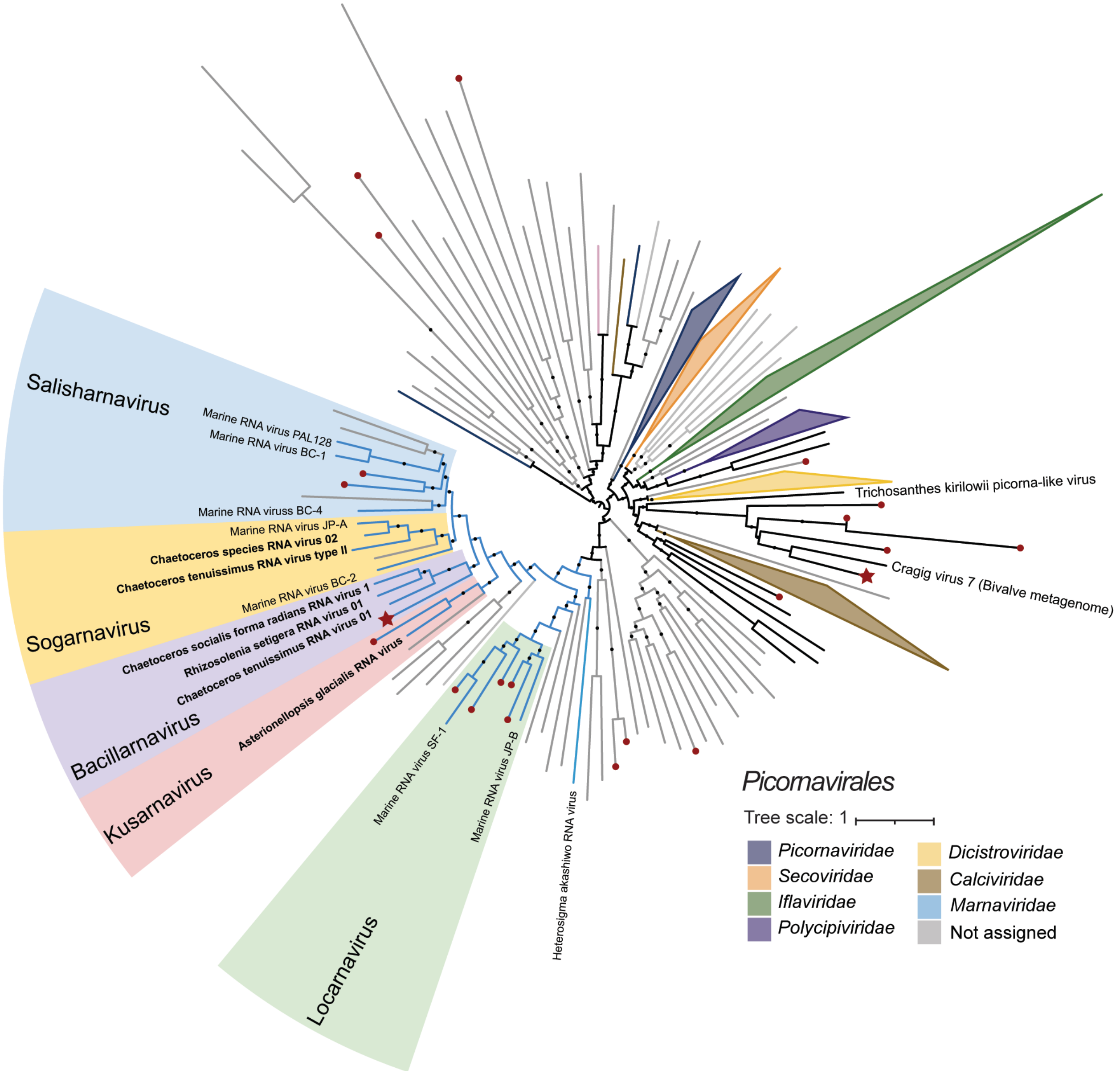

**Supplemental Figure 6.** ML phylogenetic placement of *Picornavirales* (phylum *Pisuviricota*) RdRp hallmark genes. Nodes ending in red dots represent Lake Erie RdRp. Nodes ending in red stars represent Lake Erie RdRp with greater that 0.5% contribution to average dissimilarity between the ice-covered and ice-free sample groups.

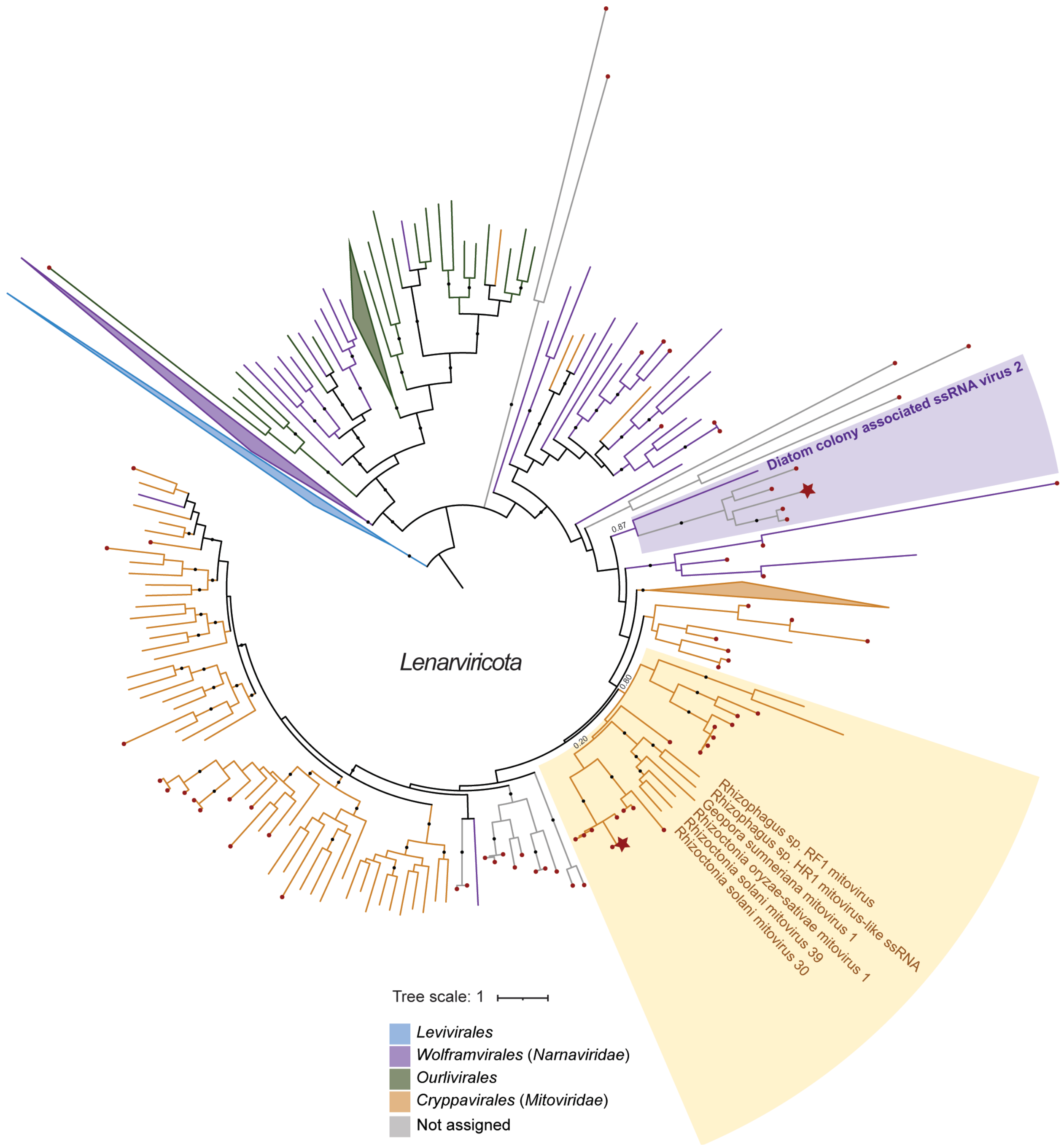

**Supplemental Figure 7.** ML phylogenetic placement of *Lenarviricota* RdRp hallmark genes. Nodes ending in red dots represent Lake Erie RdRp. Nodes ending in red stars represent Lake Erie RdRp with greater that 0.5% contribution to average dissimilarity between the ice-covered and ice-free sample groups.

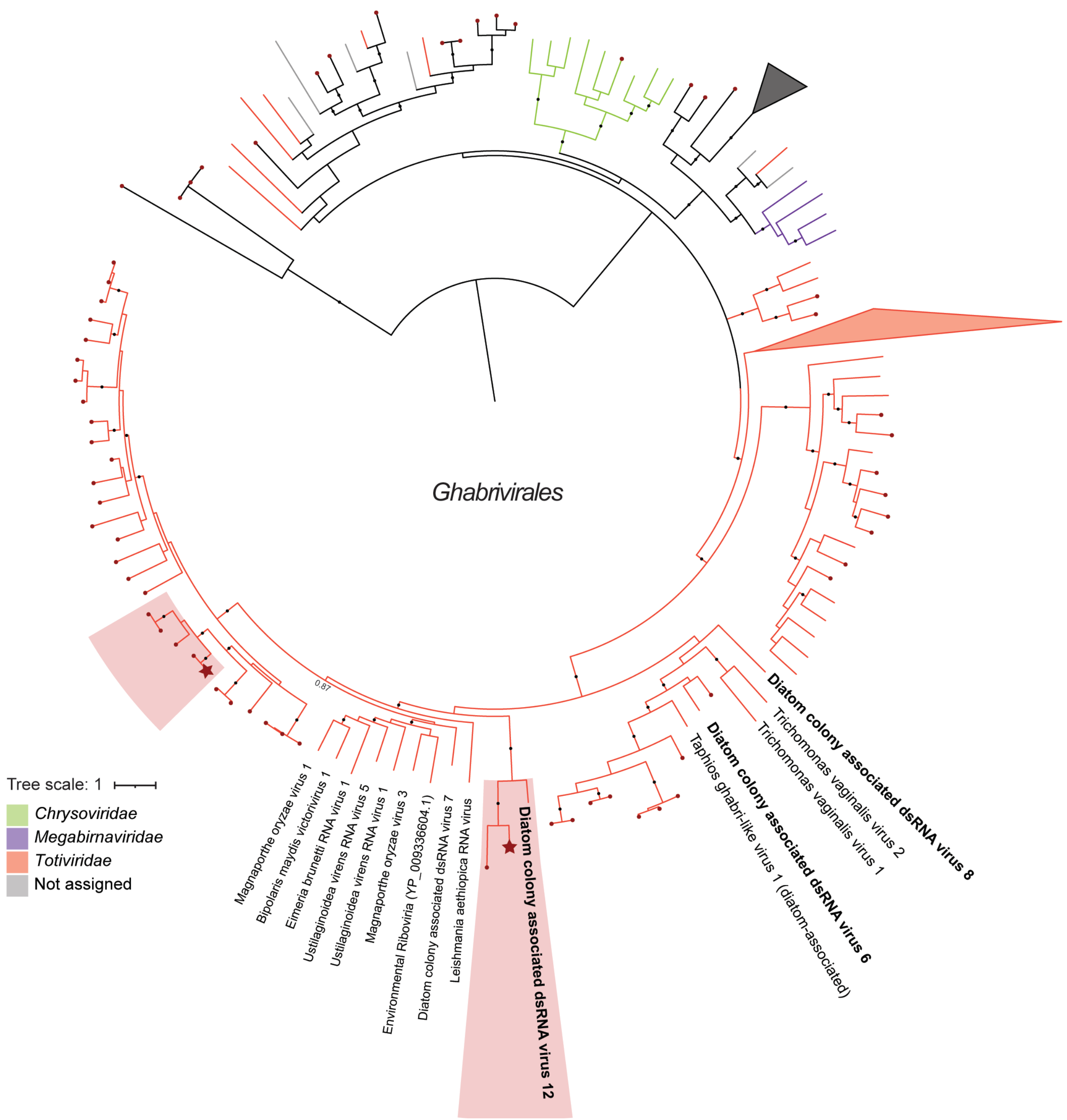

**Supplemental Figure 8.** ML phylogenetic placement of *Ghabrivirales* (phylum *Duplornaviricota*) RdRp hallmark genes. Nodes ending in red dots represent Lake Erie RdRp. Nodes ending in red stars represent Lake Erie RdRp with greater that 0.5% contribution to average dissimilarity between the ice-covered and ice-free sample groups.

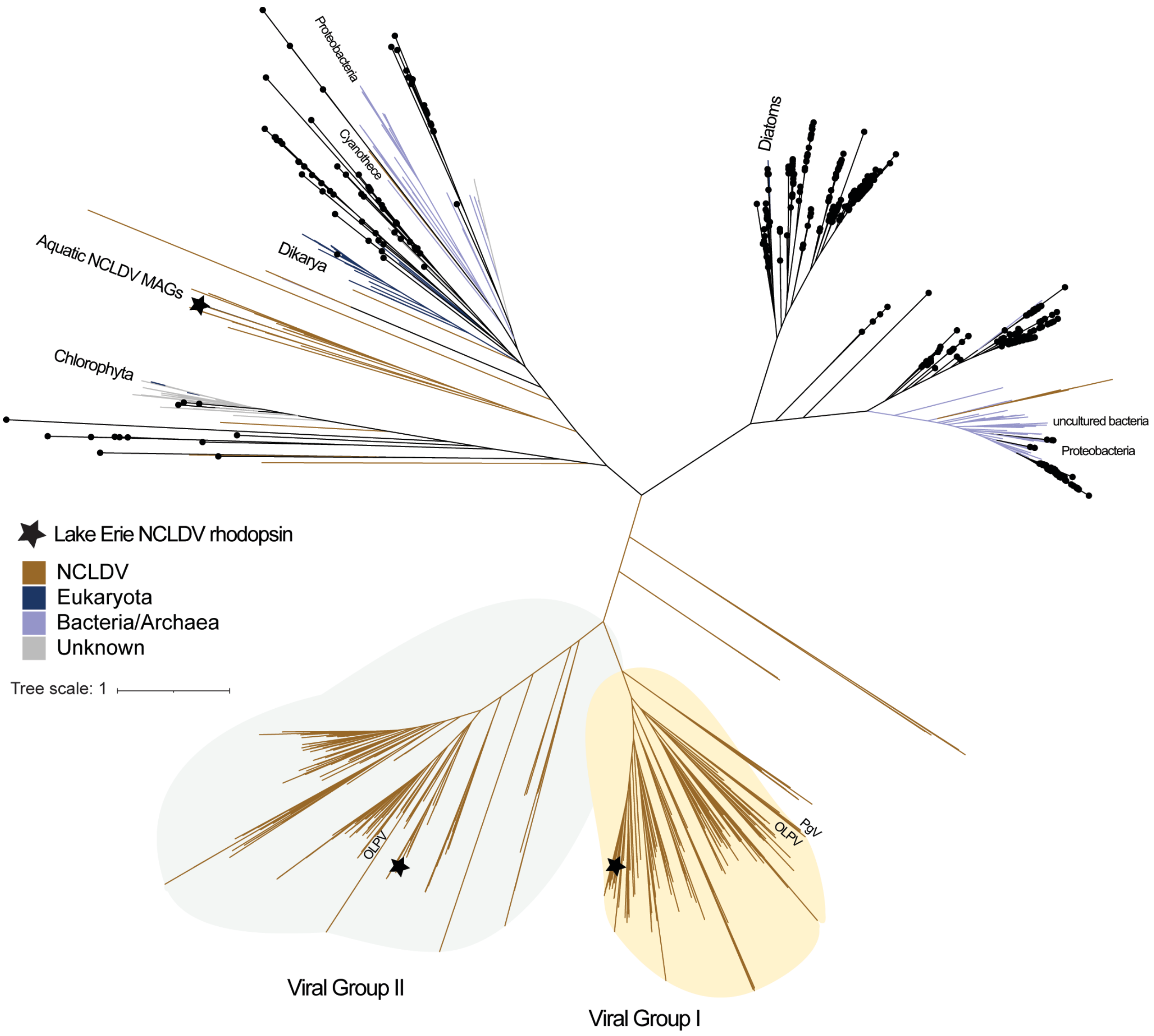

**Supplemental Figure 9.** ML phylogenetic placement of Lake Erie bacteriorhodopsin (bR). Nodes ending in black dots represent Lake Erie bR. Nodes ending in black stars represent the three putative NCLDV bR.

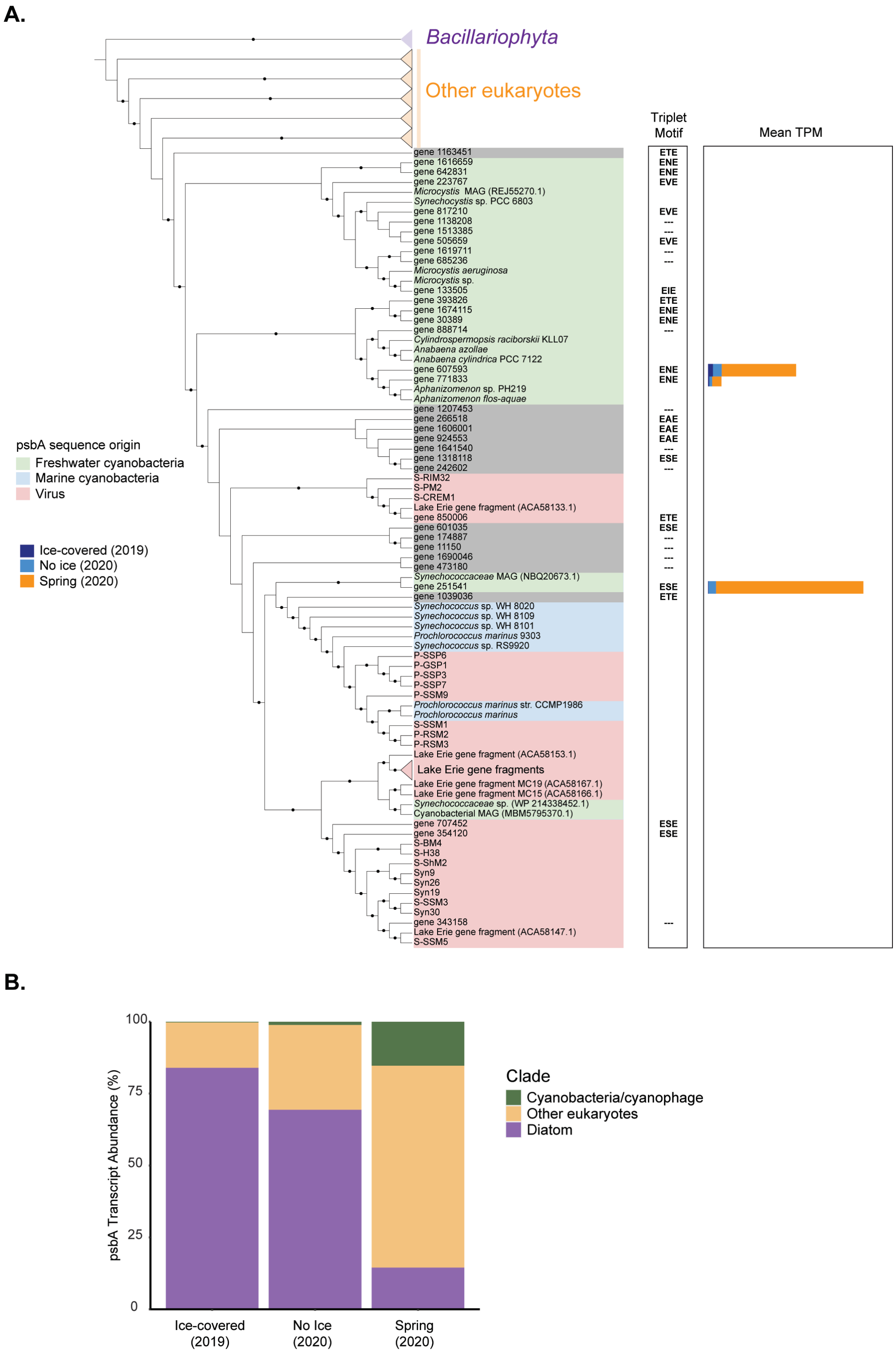

**Supplemental Figure 10. A)** ML phylogeny of PsbA sequences. PsbA >100 amino acids were used for phylogenetic analysis (342 genes). Near duplicate sequences (>99% average amino acid identity) were clustered prior to building the tree. B) Relative PsbA transcript abundance (TPM) summed by phylogenetic clade placement.

### SUPPLEMENTAL TABLES

**Supplemental Table 1**. Co-assembly statistics generated by QUAST v5.0.2. All QUAST statistics are based on contigs of size >= 500 bp unless otherwise noted.

| | |
|---|---|
| # contigs (>= 0 bp) | 1677102 |
| # contigs (>= 1000 bp) | 148925 |
| # contigs (>= 5000 bp) | 2139 |
| # contigs (>= 10000 bp) | 288 |
| # contigs (>= 25000 bp) | 8 |
| # contigs (>= 50000 bp) | 0 |
| Total length (>= 0 bp) | 966763097 |
| Total length (>= 1000 bp) | 247639904 |
| Total length (>= 5000 bp) | 15925927 |
| Total length (>= 10000 bp) | 4001501 |
| Total length (>= 25000 bp) | 258591 |
| Total length (>= 50000 bp) | 0 |
| # contigs | 639361 |
| Largest contig | 40227 |
| Total length | 574417227 |
| GC (%) | 44.46 |
| N50 | 891 |
| N75 | 641 |
| L50 | 190932 |
| L75 | 383548 |
| # N's per 100 kbp | 0 |

**Supplemental Table 2.** Viral hallmark genes identified in the metatranscriptome co-assembly *via* BLASTP.

| **Viral Group** | **Marker** | **BLASTP Database Source** | **No. Identified** |
|---|---|---|---|
| *Caudovirales* | Gp23 | PF07068 | 505 |
| | Tail Sheath | PF17481, PF17482, and PF04984 | 239 |
| | PolB (gp34) | PF03175 and PF00136 | 33 |
| | Gp20 | PF06810 | 1 |
| | Integrase | PF00589 | 1 |
| | Excisionase | PF06806 | 0 |
| | CI Repressor | PF07022 | 0 |
| | Cro Repressor | IP000655 | 0 |
| | TerL | PF05876 and PF20454 | 1 |
| *Nucleocytoviricota* | MCP | NCVOG0022 | 615 |
| | PolB | NCVOG0038 | 9 |
| *Orthornavirae* | RdRp | RdRp-Scan | 397 |
| *Cressdnaviricota* | Rep | Kazlauskas *et al.* and Moniruzzaman *et al.* | 8 |

**Supplemental Table 3.** Taxonomic estimate based on ML phylogeny of the identified viral hallmark genes. Viral hallmark gene detection in the winter (either ice-covered or ice-free samples) versus spring only was determined via read mapping.

| | | | **Detected in** | |
|---|---|---|---|---|
| | **Taxonomy estimate** | **No. hallmark genes identified** | **Winter** | **Spring only** |
| **Gp23** | Uncultured myophage | 468 | 439 | 29 |
| | Cyanomyophage | 2 | 1 | 1 |
| | *Pelagibacter* phage | 6 | 6 | 0 |
| | Uncertain | 29 | 22 | 7 |
| | **Total** | **505** | **468** | **37** |
| **MCP** | *Pimascovirales* | 24 | 10 | 14 |
| | *Asfuvirales* | 9 | 8 | 1 |
| | *Pandoravirales* | 4 | 3 | 1 |
| | *Algavirales* | 18 | 9 | 9 |
| | *Imitervirales* | 557 | 395 | 162 |
| | Uncertain | 3 | 2 | 1 |
| | **Total** | **615** | **427** | **188** |
| **RdRp** | *Pisuviricota* | 92 | 76 | 16 |
| | *Duplornaviricota* | 99 | 84 | 15 |
| | *Lenarviricota* | 84 | 72 | 12 |
| | *Kitrinoviricota* | 53 | 43 | 10 |
| | *Negarnaviricota* | 17 | 13 | 4 |
| | *Birnaviridae* | 1 | 1 | 0 |
| | *Permutotetraviridae* | 1 | 1 | 0 |
| | Uncertain | 50 | 42 | 8 |
| | **Total** | **397** | **332** | **65** |

**Supplemental Table 4.** ANOVA results for alpha diversity metrics compared between the three seasons sampled (winter 2019, winter 2020, and spring 2020). Diversity metrics were generated from the relative transcript abundance (TPM) table of all viral hallmark genes.

| | **Comparison** | **Observed richness** | **Pielou's evenness** | **Shannon's H** |
|---|---|---|---|---|
| **Main test** | **Season** | **2.20E-05** | 0.545 | 0.746 |
| **Tukey's HSD** | **Ice-covered/Ice-free** | **0.0000205** | NA | NA |
| | **Ice-free/spring** | **0.0242167** | NA | NA |
| | **Ice-covered/spring** | **0.0056772** | NA | NA |

**Supplemental Table 5.** ANOVA results for alpha diversity metrics compared between the three seasons sampled (winter 2019, winter 2020, and spring 2020). Diversity metrics were generated from the relative transcript abundance (TPM) table of separate viral hallmark gene types.

| | | **Gp23** | | |
|---|---|---|---|---|
| | **Comparison** | **Observed richness** | **Pielou's evenness** | **Shannon's H** |
| **Main test** | **Season** | **5.26E-06** | **7.36E-04** | **3.58E-02** |
| **Tukey's HSD** | **Ice-covered/Ice-free** | **1.93E-05** | 7.42E-02 | 9.78E-01 |
| | **Ice-free/spring** | 3.50E-01 | 2.78E-01 | 1.37E-01 |
| | **Ice-covered/spring** | **1.03E-04** | **5.91E-04** | **3.42E-02** |

| | | **MCP** | | |
|---|---|---|---|---|
| | **Comparison** | **Observed richness** | **Pielou's evenness** | **Shannon's H** |
| **Main test** | **Season** | **1.38E-02** | **2.50E-02** | **1.25E-03** |
| **Tukey's HSD** | **Ice-covered/Ice-free** | **3.12E-02** | 7.27E-01 | **9.04E-04** |
| | **Ice-free/spring** | **1.37E-02** | 2.27E-01 | **1.17E-02** |
| | **Ice-covered/spring** | 7.12E-01 | **1.96E-02** | 5.63E-01 |

| | | **RdRp** | | |
|---|---|---|---|---|
| | **Comparison** | **Observed richness** | **Pielou's evenness** | **Shannon's H** |
| **Main test** | **Season** | **2.73E-05** | 1.09E-01 | **3.53E-02** |
| **Tukey's HSD** | **Ice-covered/Ice-free** | **1.16E-04** | NA | 4.34E-01 |
| | **Ice-free/spring** | 5.80E-01 | NA | 5.18E-01 |
| | **Ice-covered/spring** | **3.03E-04** | NA | **2.89E-02** |

**Supplemental Table 6.** Spearman correlations between various diversity and abundance metrics. Significant correlations are bolded and shown in grey.

| | | **Spearman** | |
|---|---|---|---|
| **Comparison** | | **rho** | **p** |
| ***Bacillariophyta* TPM** | **Prokaryotic TPM** | -0.8329670 | **0.0002166** |
| ***Bacillariophyta* TPM** | **RpoB TPM** | -0.7098901 | **0.004451** |
| *Bacillariophyta* TPM | RpoB richness | -0.2615385 | 0.3664 |
| *Bacillariophyta* TPM | Rpb1 richness | -0.0044249 | 0.988 |
| *Bacillariophyta* TPM | RpoB TPM | 0.1868132 | 0.5225 |
| *Bacillariophyta* TPM | Rpb1 TPM | -0.2615385 | 0.3664 |
| **Prokaryotic TPM** | **Viral hallmark gene richness** | 0.9384615 | **6.86E-07** |
| ***Bacillariophyta* TPM** | **Viral hallmark gene richness** | -0.8153846 | **0.0003791** |
| ***Bacillariophyta* TPM** | **Gp23 TPM** | -0.7318681 | **0.002925** |
| *Bacillariophyta* TPM | MCP TPM | -0.1340659 | 0.6477 |
| ***Bacillariophyta* TPM** | **RdRp TPM** | -0.8769231 | **3.83E-05** |
| *Bacillariophyta* TPM | Viral hallmark gene richness | -0.1956044 | 0.5028 |
| *Bacillariophyta* TPM | Gp23 richness | -0.1870188 | 0.522 |
| *Bacillariophyta* TPM | MCP richness | -0.1584159 | 0.5886 |
| ***Bacillariophyta* TPM** | **RdRp richness** | -0.6072611 | **0.02127** |
| Prokaryotic TPM | Gp23 richness | 0.4576460 | 0.09988 |
| **Prokaryotic TPM** | **MCP richness** | 0.5808584 | **0.02939** |
| **Prokaryotic TPM** | **RdRp richness** | 0.6226626 | **0.01739** |
| **RpoB/Rpb1 richness** | **Viral hallmark gene richness** | 0.9340659 | **1.028E-06** |
| **RpoB richness** | **Gp23 richness** | 0.9636970 | **3.06E-08** |
| **RpoB richness** | **MCP richness** | 0.7876792 | **8.23E-04** |
| **RpoB richness** | **RdRp richness** | 0.6600664 | **1.02E-02** |
| **Rpb1 richness** | **MCP richness** | 0.7497346 | **0.002018** |
| Rpb1 richness | RdRp richness | 0.4507267 | 0.1058 |
| **Rpb1 richness** | **Gp23 richness** | 0.8815196 | **3.08E-05** |

**Supplemental Table 7.** Select output from similarity percentage (SIMPER) analysis. Viral hallmark genes contributing to the top ~10% cumulative dissimilarity between the ice-covered and ice-free winter are shown (see Figure 4). The abundance values shown are based on the square root transformed TPM table.

| | | **Ice-covered (2019)** | **Ice-free (2020)** | | | | |
|---|---|---|---|---|---|---|---|
| **Gene ID** | **Viral marker** | **Av. Abund.** | **Av. Abund.** | **Av. Diss.** | **Diss/SD** | **Contrib. (%)** | **Cumul. (%)** |
| gene_458515 | RdRp | 1.94 | 22.95 | 1.46 | 2.25 | 2.06 | 2.06 |
| gene_293376 | Gp23 | 8.46 | 24.46 | 1.2 | 1.65 | 1.7 | 3.76 |
| gene_678216 | Gp23 | 4.87 | 15.75 | 0.81 | 1.71 | 1.14 | 4.9 |
| gene_1385399 | RdRp | 0 | 9.83 | 0.68 | 3.12 | 0.97 | 5.86 |
| gene_26037 | RdRp | 9.23 | 0.87 | 0.62 | 1.07 | 0.87 | 6.73 |
| gene_1263456 | RdRp | 2.68 | 8.71 | 0.43 | 2.3 | 0.61 | 7.35 |
| gene_430917 | RdRp | 10.14 | 13.89 | 0.41 | 1.19 | 0.58 | 7.93 |
| gene_1656192 | Gp23 | 5.26 | 10.08 | 0.41 | 1.64 | 0.58 | 8.51 |
| gene_626655 | Gp23 | 0 | 5.56 | 0.39 | 2.7 | 0.56 | 9.06 |
| gene_1534634 | RdRp | 6.68 | 2.38 | 0.38 | 1.25 | 0.53 | 9.6 |
| gene_118358 | Gp23 | 8.14 | 3.73 | 0.37 | 1.81 | 0.52 | 10.12 |
